# Upregulation of steroidogenesis is associated with coma in human cerebral malaria

**DOI:** 10.1101/2023.05.01.538900

**Authors:** Wael Abdrabou, Himanshu Gupta, Praveen K. Sahu, Akshaya K. Mohanty, Rajyabardhan Pattnaik, Angelika Hoffmann, Sanjib Mohanty, Samuel C. Wassmer, Youssef Idaghdour

## Abstract

*P. falciparum* parasites manipulate host metabolic processes during malaria to ensure their survival and progression, causing the host and parasite metabolic pathways to become intertwined. We analyzed metabolites to evaluate their potential as biomarkers for cerebral malaria (CM). Our analysis of CM (with coma) and CM-like (without coma) patients identified 835 metabolites, including lipids, amino acids, xenobiotics, peptides, nucleotides, carbohydrates, cofactors, and vitamins. Principal component analysis revealed clear segregation between CM-like and CM patients. Metabolite-by-metabolite analysis identified 103 differentially abundant metabolites, 26 of which were significantly lower in CM-like patients (primarily lipids), while 71% of those higher in CM patients were amino acids and xenobiotics. The results revealed significant differences in circulating levels of long chain free fatty acids and catecholamine metabolism and identified steroid biosynthesis as the most enriched lipid metabolism pathway, with eight endogenous steroids showing significantly higher levels in CM patients compared to CM-like patients (FC > 2, B-H FDR-adjusted P < 0.05). These steroids include pregnenolone sulfate, pregnenediol sulfate, pregnenetriol sulfate, androsterone monosulfate, 16α-OH-DHEA-S, DHEA-S, cortisol, and cortisone. High levels of pregnenolone and its downstream metabolic derivatives were significantly associated with coma in CM patients. Our findings suggest that monitoring circulating neurosteroid levels in patients could aid in the early identification of those at risk of coma, and may important implications for the clinical management of CM patients.

## INTRODUCTION

*Plasmodium falciparum* infection is a major global health concern, with an estimated 247 million clinical cases and 619,000 deaths in 2021^1^. Cerebral Malaria (CM) is a severe complication of *falciparum* malaria consisting of an acute encephalopathy characterized by coma and a mortality rate up to 25%^2,3^. The etiology of encephalopathy is not well understood but likely to be multifactorial. Coma is a state of prolonged unconsciousness in which a person does not respond to stimuli. It can be caused by a variety of factors, including traumatic brain injury (TBI), stroke, drug overdose, and infections. The pathophysiology of coma is not fully understood, but it is thought to involve alterations in neuronal excitability and synaptic transmission^4^. Treatment for coma typically involves addressing the underlying cause and providing supportive care, such as oxygen therapy, intravenous fluids, and nutrition^5^.

There have been limited investigations aimed at gaining a deeper understanding of the cause of coma in CM. An autopsy study in Vietnamese adults with CM demonstrated that edema, plasma protein leakage, and vascular endothelial growth factor signaling did not correlate with pre-mortem coma^6^. More recently, an abnormal elevation of the neuro-modulatory amino acid pipecolic acid (PA) was reported in the plasma of Malawian children with CM, and parasite-derived PA was linked to neurological impairment in the murine experimental model of CM, suggesting a role in the development of coma^7^. Our group used Apparent Diffusion Coefficient (ADC), a measure of the magnitude of diffusion of water molecules within tissues and commonly clinically calculated using magnetic resonance imaging (MRI) with diffusion-weighted imaging, to investigate the cause of death in adult CM^8^. We showed that reversible, hypoxia-induced cytotoxic edema identified by decreased ADC signal occurred predominantly in the basal ganglia of adults with non-fatal CM^9^. Remarkably, we also found that ∼20% of patients with severe, non-cerebral malaria (SNCM) presented similar brain alterations despite the absence of deep coma^9^.

Here we investigate metabolic changes associated with coma in an Indian cohort^8,9^. To achieve this goal, we conducted global untargeted metabolomic profiling of plasma samples from a subset of patients with CM (coma and decreased ADC signal in the basal ganglia) and CM-like (SNCM without coma, decreased ADC signal in the basal ganglia) to identify metabolites and metabolic processes specifically linked to a deep state of unconsciousness.

## METHODS

### Sample collection

Samples were collected as part of a parent study^8,9^. Patients with *P. falciparum* malaria were admitted to Ispat General Hospital in Rourkela, India, from October 2013 to November 2019. CM and SNCM adult patients were selected for this analysis using the modified World Health Organization (WHO) criteria^3^, and the Glasgow Coma Score (GCS) was used to assess levels of consciousness. CM patients (n=6) were defined by a GCS <11 out of 15 after correction of hypoglycemia (<2.2 mmol/l), presented asexual forms of *P. falciparum* in a peripheral blood smear.

CM-like patients (n=4) were defined by a GCS 11-15 with one or more complications including severe malarial anemia (Hb <7 g/dL, n=0/4), jaundice (bilirubin >3 mg/dL, n=3/4), acute kidney injury (serum creatinine >3 mg/dL, n=3/4), and hyperlactatemia (lactate >5 mmol/L, n=0/4); they were selected from a larger group of SNCM patients due to their CM-like MRI pattern^9^. (**Fig. 1a**). Patient characteristics are presented in **Table 1**; they were all treated according to Indian national guidelines^10^. Exclusion criteria included other plasmodial species co-infection, meningitis, or other causes of encephalopathy, as well as bacterial infection, as detailed elsewhere^11^. We used consecutive sampling for this study, a non-probability sampling technique that seeks to include all accessible subjects as part of the sample. Because Ispat General Hospital is the only referral health facility for severe *falciparum* malaria in Rourkela and its adjoining districts, this approach ensured that the sampling during the 6 years of enrolment was highly representative of the clinical cases in this locality. Ethical approval was obtained from the Indian Council of Medical Research (TDR589/2010/ECDII), as well as from the institutional review boards from New York University School of Medicine (S12-03016), the London School of Hygiene and Tropical Medicine, and Ispat General Hospital. A signed informed consent was obtained from all participants and/or their legal guardians. In accordance with the Health Insurance Portability and Accountability Act, patient details were kept confidential using unique study numbers.

**Table 1:**
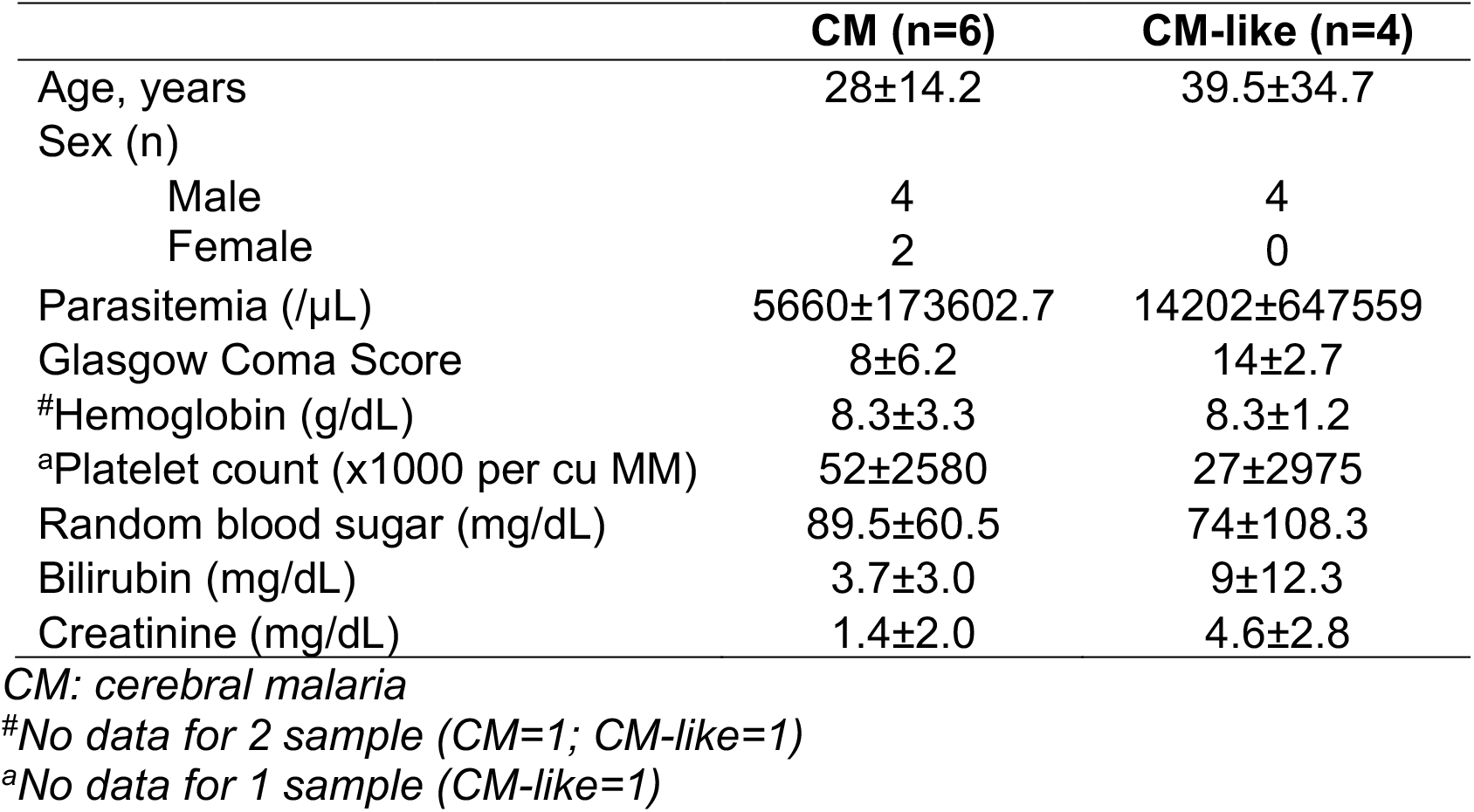
Characteristics of Indian patients with CM and CM-like malaria recruited from October 2013 to November 2019. Values are expressed in median± interquartile range.

**Fig. 1.**
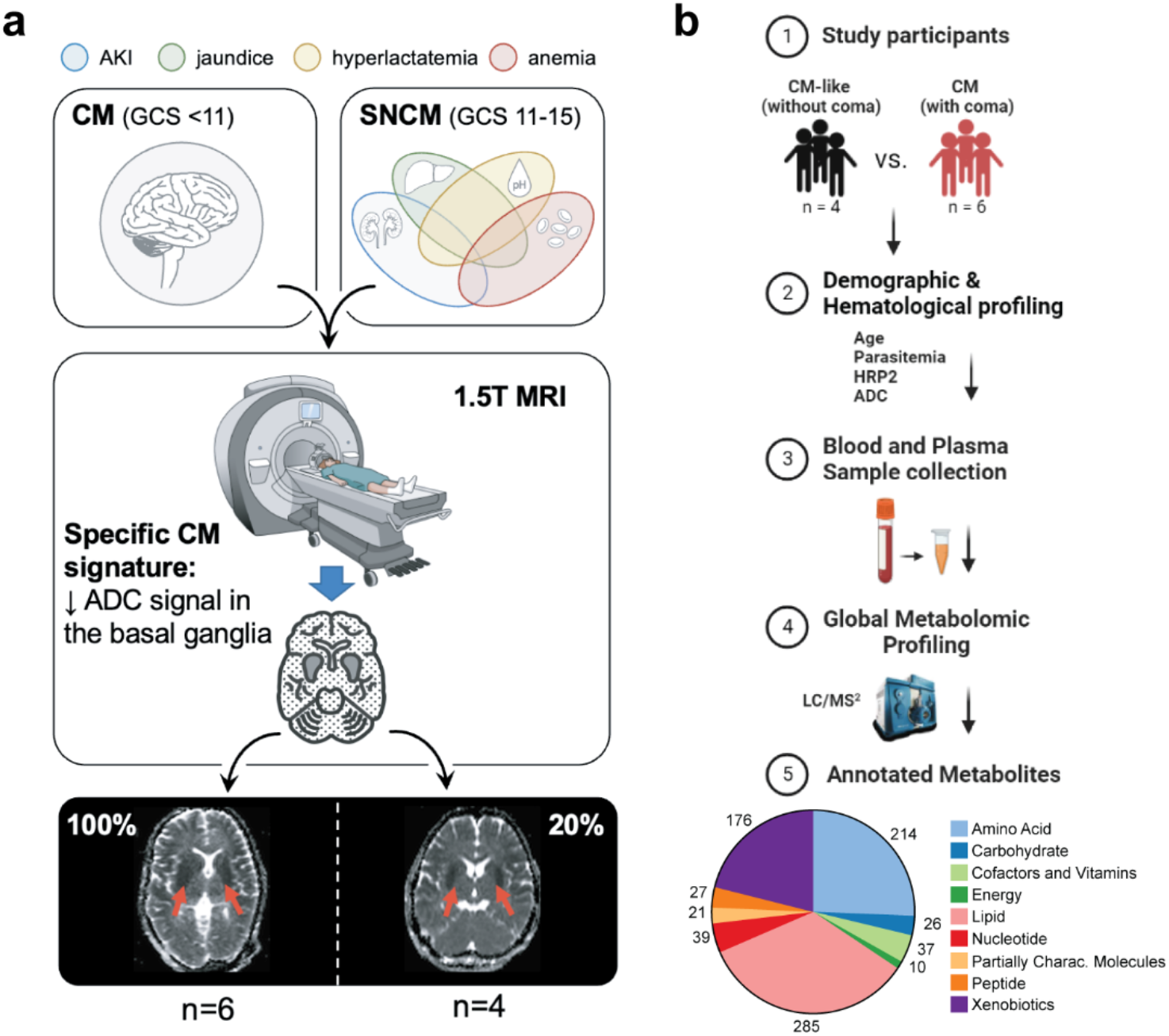
Study design and metabolomic profiling of CM patients with and without coma. **a**, An illustration showing patient selection criteria. *P. falciparum*-infected cerebral malaria (CM) and severe non-cerebral malaria (SNCM) adult patients were selected using the modified World Health Organization (WHO) criteria, and the specific CM signature of decreased Apparent Diffusion Coefficient (ADC) in basal ganglia. The Glasgow Coma Score (GCS) was used to assess levels of consciousness. CM patients (with coma, n = 6) were defined by a GCS <11 out of 15. CM-like patients (without coma, n = 4) were defined by a GCS 11-15. **b**, A schematic showing the study design, sample size, and the outcome of global untargeted metabolomic profiling of plasma samples collected from CM and CM-like patients. Total number of metabolites identified using LC/MS^2^ and retained for downstream analysis is shown (n = 835). Each color represents a metabolite class. The number of metabolites for each class is shown.

### Global untargeted metabolomic profiling

Blood samples were collected and plasma isolated using standard protocols. A volume of 100 μl of plasma from the 6 CM and 4 CM-like patients were shipped in dry ice from India to Metabolon for global untargeted metabolomic profiling. Following receipt, samples were inventoried and immediately stored at -80C. Samples were prepared using the automated MicroLab STAR® system (Hamilton) for metabolite profiling. Several recovery standards were added prior to the first step in the extraction process for quality control purposes. To remove protein, dissociate small molecules bound to protein or trapped in the precipitated protein matrix, and to recover chemically diverse metabolites, proteins were precipitated with methanol under vigorous shaking for 2 min (Glen Mills GenoGrinder 2000) followed by centrifugation. The resulting extract was divided into five fractions. Two of these fractions were subjected to analysis using two distinct reverse phase (RP)/UPLC-MS/MS methods, both utilizing positive ion mode electrospray ionization (ESI). Additionally, one fraction was analyzed through RP/UPLC-MS/MS with negative ion mode ESI, another through HILIC/UPLC-MS/MS with negative ion mode ESI, and a final fraction was set aside as a backup. Samples were placed briefly on a TurboVap® (Zymark) to remove the organic solvent. The sample extracts were stored overnight under nitrogen before preparation for analysis described elsewhere.

### Metabolomic data curation

The raw data was extracted and analyzed to identify signature chromatographic peaks and relative ion concentrations for each detected metabolite in the plasma samples. To identify and quantify individual components, we used the Quantify Individual Components in a Sample (QUICS) method^12^ to analyze the spectrometry data. The QUICS method groups ions for any given metabolite from LC-MS/MS or GC-MS data based on retention time, mass, ion intensity, and covariance of the ion data across the overall set of samples^12^. To perform metabolite identification, each metabolite aggregate was matched to a thoroughly annotated reference chemical library comprising over 4000 metabolites with well-defined chemical profiles, including the retention time/index (RI), mass-to-charge ratio (m/z), and chromatographic data, including MS/MS spectral data^12–14^. Chromatographic peaks were quantified using the area-under-the-curve method, and the normalized values of each metabolite for all participants are provided in **Supplementary Table 1**. Data normalization was performed to remove variation resulting from instrument inter-run tuning differences, where each compound was corrected in run blocks by registering the medians to equal one (1.00) and normalizing each data point proportionately. Next, we imputed missing values with the observed minimum after normalization.

### Statistical analyses

Differences in age, ADC, parasitemia count, parasite biomass by HRP2 and endothelial dysfunction biomarker such as angiopoietin between CM-like and CM patients were evaluated by performing two-tailed unpaired Mann–Whitney tests using GraphPad Prism v8.0. The curated data for each detected metabolite were log-transformed, and interquartile (IQR) normalized using SAS/JMP Genomics version 8.0 (SAS Institute Inc., Cary, NC) to eliminate potential technical artifacts and outliers. Unsupervised statistical analyses, including principal component analysis (PCA) was conducted to investigate the correlation structure in the data and global influences on the metabolome across the two groups. All supervised statistical analyses were performed using SAS/JMP Genomics version 8.0 (SAS Institute). Analysis of covariance (ANCOVA) was utilized to identify differentially abundant metabolites that were statistically significant between the two groups. A false discovery rate threshold of 0.05 (Benjamini-Hochberg FDR) was used to determine statistical significance.

### Quantitative pathway enrichment analysis

The Human Metabolome Database (HMDB, www.hmdb.ca)^15^ was used to query the mass of each detected metabolite. The differentially abundant metabolites annotated by the HMDB database and identified by supervised statistical analyses^15^ the Kyoto Encyclopedia of Genes and Genomes (KEGG) and subjected to quantitative enrichment analysis (QEA) using the online software MetaboAnalyst 5.0 (http://www.metaboanalyst.ca/)^16^. Pathway enrichment analysis was performed using the *globaltest* package^17^, which employs a generalized linear model to estimate a Q-statistic for each metabolite set, reflecting the correlation between compound concentration profiles and clinical outcomes. The Q-statistic for a metabolite set is the average of the Q-statistics for each metabolite in the set. Significantly enriched metabolic pathways were defined as those with FDR-adjusted *<u>p</u>*-values < 0.05.

## RESULTS AND DISCUSSION

### Characteristics of the study participants

The levels of consciousness of participating patients were assessed using the GCS. CM patients (with coma, n=6) were defined as having a GCS score of less than 11 out of 15 after correction of hypoglycemia, with the presence of asexual forms of *P. falciparum* in a peripheral blood smear. CM-like patients (without coma, n=4) were defined as having a GCS score of 11-15 and one or more complications, such as severe malarial anemia, jaundice, acute kidney injury, and hyperlactatemia (**Fig. 1a**). Importantly, both groups were matched for age and levels of parasitemia. Additionally, no significant difference was observed between the two groups in average parasitic biomass, ADC or levels of angiopoietin-2 (**Fig. 2**).

**Fig. 2.**
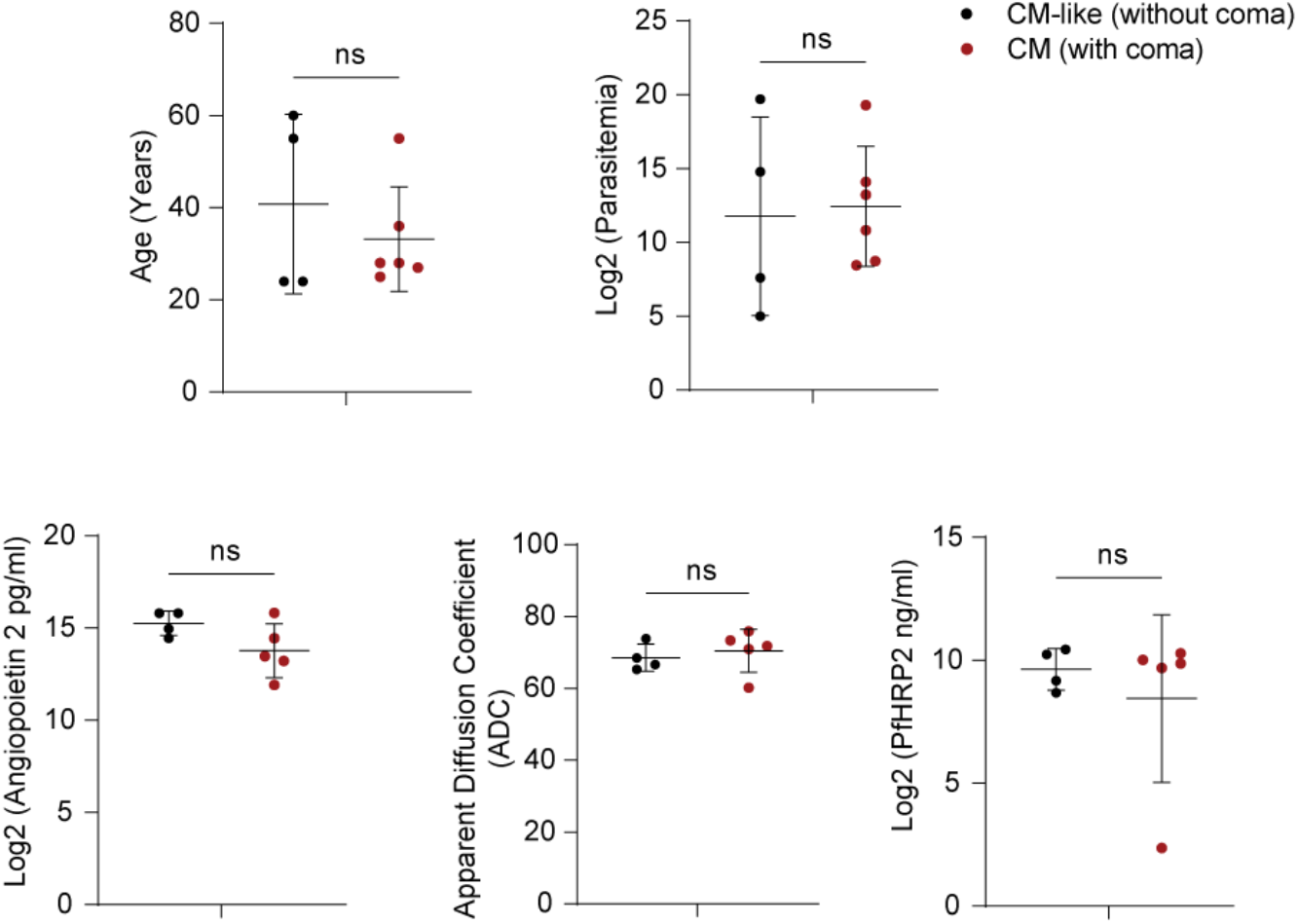
Study groups patient characteristics comparisons. Scatter dot plots showing differences between the two study groups CM-like (n = 4, black) and CM (n = 6, red) in average age (years) log2 parasitemia, parasitic biomass (Log2 PfHRP2), endothelial dysfunction biomarker (Log2 angiopoietin-2 pg/ml) and Apparent Diffusion Coefficient (ADC). Statistical significance in the comparisons shown was assessed using a paired two-tailed Student’s *t*-test (*P* = ns: non-significant). Bar and whiskers represent mean ± SD.

### Global untargeted metabolomic profiling

Stringent quality control of the global metabolomic data led to the retention of 835 annotated metabolites for downstream analysis. Most detected metabolites were lipids (n = 285) and amino acids (n = 214). The third most represented subgroup consisted of xenobiotics (n = 176, **Fig. 1b**). The majority of the xenobiotics detected were involved in benzoate and xanthine metabolism, exogenous food/plant components, or were components of generic analgesics and antibiotics. In addition, the metabolites detected included 27 peptides, 39 nucleotides, 26 carbohydrates, 37 cofactors and vitamins, 10 energy metabolites, and 21 partially characterized molecules (**Fig. 1b**). PCA of the full dataset revealed a clear segregation of CM-like and CM patients, with the first two principal components explaining 42.8% of the variation in the dataset (**Fig. 3a**). Next, metabolite-by-metabolite analysis (**Fig. 3b**) revealed 103 differentially abundant metabolites (FC ≥ |1.5|, B-H FDR < 0.05, **Fig. 3c**), 26 of which showed significantly lower levels in CM-like patients relative to CM patients. Remarkably, 85% of these metabolites were lipids, 14 of which were long-chain fatty acids and eight steroids (**Fig. 3c** and **Supplementary Table 2**). In contrast, 71% of the metabolites that showed significantly higher levels in CM patients relative to CM-like patients were amino acids (n = 42) and xenobiotics (n =13) (**Fig. 3c** and **Supplementary Table 2**). Together, these results show that CM-like and CM patients have distinct metabolic signatures.

**Fig. 3:**
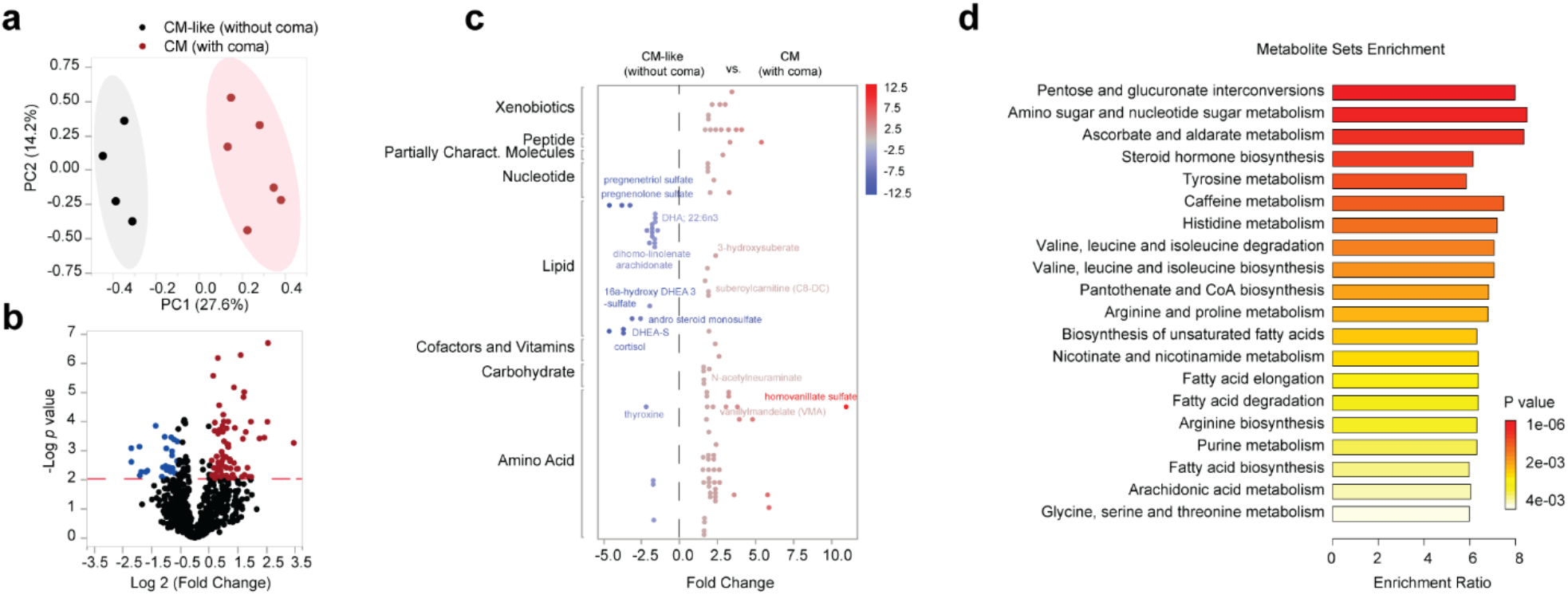
Metabolic differences between CM patients with and without coma. **a**, PCA of the global metabolome of CM-like (n = 4, black) and CM (n = 6, red) patients. **b**, Volcano plot of differential metabolite abundance (n = 835 metabolites). Significant differentially abundant metabolites between the two study groups (repeated measures ANCOVA, B-H FDR < 0.05, log2 |Fold Change| ≥ 1.5) are shown in red (n = 77) and blue (n = 26). Significantly upregulated (red) and downregulated (blue) metabolites in CM-like patients relative to CM patients are shown to the right and left of the volcano plots, respectively. **c**, Classes of differentially abundant metabolites between CM-like and CM patients are shown (n = 103, B-H FDR < 0.05, log2 |Fold Change| ≥ 1.5). Metabolites are colored (blue to gray to red) to indicate their fold change from significantly low to significantly high. **a**, Metabolite set enrichment analysis. Results of the enrichment analysis of 103 differentially abundant metabolites using MetaboAnalyst v5.0. Horizontal bars represent pathways’ fold enrichment and the color gradient indicate statistical significance from low (yellow) to high (red). Pathways are ranked from top to bottom by significance (*P* value).

### Metabolic pathway enrichment analysis

Quantitative enrichment analysis (QEA) of the 103 differentially abundant metabolites using MetaboAnalyst 5.0^16^ revealed 21 significantly enriched metabolic KEGG pathways that exhibited distinct differences between CM-like and CM patients (**Fig. 3d** and **Supplementary Table 3**). Among the enriched pathways, the top three most significantly enriched pathways were pentose and glucuronate interconversions, amino and nucleotide sugar metabolism, and ascorbate and aldarate metabolism. These pathways are involved in the use and interconversion of common metabolites such as glucose, glucuronate, galacturonate, xylose, glucosamine, galactosamine, and nucleotide sugars (B-H FDR-adjusted *P* < 4.54e-05; **Fig. 3d**). Notably, the plasma levels of gulonic acid, L-arabitol, and N-acetylneuraminic acid (NeuAc) implicated in these pathways were significantly elevated in CM-like patients (FC ≥ |1.5|, B-H FDR-adjusted *P* <0.05, **Supplementary Table 2 and 3**). NeuAc is an indispensable part of the important functional sugars, sialic acids, which play a critical role in maintaining and improving brain health. The plasma levels of NeuAc have been found to correlate with brain wet weight and brain ganglioside and glycoprotein NeuAc^18^.

Of the enriched pathways, amino acid and lipid metabolism pathways, specifically steroid biosynthesis and fatty acid metabolism, were the most represented. The most significantly enriched amino acid metabolism pathway was tyrosine metabolism (B-H FDR-adjusted *P* = 5.18e-4), followed by histidine metabolism (B-H FDR-adjusted *P* = 1.16e-3), and valine, leucine, and isoleucine degradation and biosynthesis (B-H FDR-adjusted *P* = 1.16e-3). The most significantly enriched lipid metabolism pathway was steroid biosynthesis (B-H FDR-adjusted *P* = 2.70e-4), and fatty acid metabolism pathways were also represented (e.g., biosynthesis of unsaturated fatty acids (B-H FDR-adjusted *P* = 5.18.e-4)) (**Fig. 3d**). **Supplementary Table 3** provides a full list of the enriched metabolic pathways. The results of the QEA analysis demonstrate that the metabolic signatures of CM-like and CM patients differ significantly in various metabolic pathways, implicating processes and metabolites that are critical for brain health.

### Differential circulating levels of free fatty acids and acylcarnitines

QEA analysis revealed a significant over-representation of multiple fatty acid metabolic pathways, including biosynthesis of unsaturated fatty acids, fatty acid biosynthesis, fatty acid elongation, fatty acid degradation, and arachidonic acid metabolism (**Fig. 3d**). Our results show that CM patients have significantly lower plasma levels of branched, hydroxyl, and dicarboxyl short- and medium-chain acylcarnitines (3-hydroxysuberate, 2-methylmalonylcarnitine (C4-DC), 3-hydroxyoctanoylcarnitine, suberoylcarnitine (C8-DC), and pimeloylcarnitine/3-methyladipoylcarnitine (C7-DC)) (FC < -1.5; B-H FDR-adjusted *P* < 0.05, **Fig. 4a**). Acylcarnitines are formed through the esterification of fatty acids (i.e., acyl groups) and L-carnitine, which is essential for shuttling fatty acids between the cytosol and mitochondria where they are used in β oxidation for energy production^19^. Carnitine shuttle is also used to transfer dicarboxylic fatty acids, which are the preferential fatty acids for peroxisomal fatty acid β oxidation^19^. The higher levels of dicarboxylic medium-chain acylcarnitines such as suberoylcarnitine (C8-DC) and 3-methyladipoylcarnitine (C7-DC) in CM-like patients are indicative of increased α- and ω-oxidation of omega-3 polyunsaturated fatty acids, as well as peroxisomal FA β oxidation^20^.

**Fig. 4:**
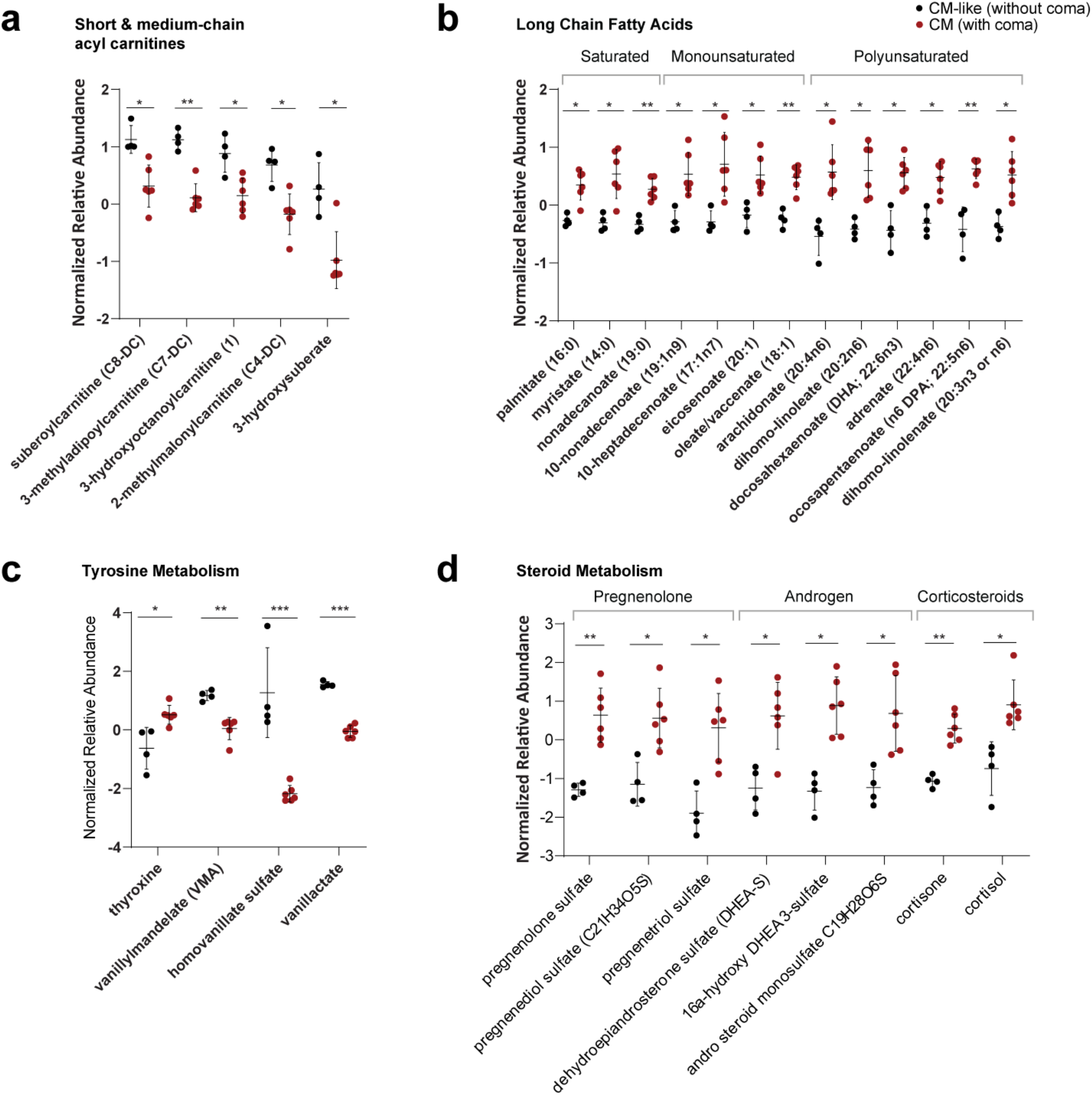
Signature fatty acid, tyrosine and neurosteroids metabolic differences. Scatter dot plots displaying the difference in normalized relative abundance levels of **a**, short and medium-chain acylcarnitines (n = 5) **b**, long-chain free fatty acids (FFAs) (n = 13) **c**, tyrosine metabolites (n = 4) between CM-like (n = 4, black) and CM patients (n = 6, red). Bar and whiskers represent mean ± SD. Statistical significance was assessed using repeated measures ANCOVA (B-H FDR-adjusted *p* value < 0.05, log2 |Fold Change| ≥ 1.5) (\**p* < 0.05 and \*\**p* < 0.01).

Conversely, we show that CM patients have significantly higher levels of 13 circulating long chain free fatty acids (FFAs) relative to CM-like patients (FC > 1.5, B-H FDR-adjusted *P* < 0.05) (**Fig. 4b**). These long-chain fatty acids include polyunsaturated (arachidonate (20:4n6), docosapentaenoate (n6 DPA; 22:5n6), dihomo-linoleate (20:2n6), dihomo-linolenate (20:3n3 or n6), and adrenate (22:4n6)); monounsaturated (10-heptadecenoate (17:1n7), 10-nonadecenoate (19:1n9), oleate/vaccenate (18:1), and eicosenoate (20:1)); and saturated (myristate (14:0) and palmitate (16:0)) fatty acids (**Fig. 4b** and **Supplementary Table 2**). FFAs in the bloodstream primarily originate from the lipolysis of triacylglycerol, which is stored as lipid droplets by the activity of classic lipases^21^. During periods of high energy demands, such as acute cold exposure, exercise, and hypoglycemia resulting from fasting or an increase in adrenergic stimulation, FFAs are mobilized and utilized in β-oxidation for energy production^22^. Our findings suggest that CM patients experience increased lipolysis relative to CM-like patients, however, it also hints to decreased mobilization and uptake of FFAs. Patients with β-oxidation defects may experience coma and brain edema as toxic intermediates accumulate^23^. In addition, blood-brain barrier dysfunction may also contribute to coma and brain edema in these patients^23^.

### Catecholamine degradation and tyrosine metabolism

QEA analysis also identified the tyrosine metabolic pathway as the most significantly enriched amino acid metabolic pathway (**Fig. 3d**). This finding is of particular interest given the direct involvement of tyrosine in the synthesis of thyroid hormones and catecholamines and their link to fatty acid metabolism and brain function. Tyrosine is the main precursor for endocrine thyroid hormones, thyroxine (T4) and triiodothyronine (T3), which are synthesized through the iodination and coupling of tyrosine residues in thyroglobulin of the thyroid gland^24^. Both hormones play an important role in regulating different biological functions from metabolic rate to cerebral maturation. Tyrosine is also the primary precursor for the synthesis of catecholamines (e.g., dopamine, norepinephrine, and epinephrine), neuromodulatory hormones that play critical role in determining the general level of metabolic activity^22^. Tyrosine is converted to L-Dopa and then to dopamine through the actions of the enzymes tyrosine hydroxylase (TH) and aromatic amino acid decarboxylase (AADC), respectively^25^. Dopamine is then converted to norepinephrine and epinephrine^22^.

Our data show that CM patients have significantly higher levels of thyroxine relative to CM-like patients (FC = 2.2, B-H FDR-adjusted *P* = 4.55e-2) (**Fig. 4c**). Hyperthyroidism can result in a thyroid hormone-mediated increase in lipolysis, circulating free fatty acids (FFAs) concentration, and mitochondrial β-oxidation of fatty acids^26,27^. Thyroid hormones stimulate lipolysis from fat stores in white adipose tissue and from dietary fat sources to generate FFAs^28^. Thyroid hormones also regulate the activity of the two main cytosolic lipases, hepatic lipase and adipose triglyceride lipase (ATGL), and transcriptionally stimulate carnitine O-palmitoyltransferase 1 (CPT1-Lα), the rate-limiting enzyme for mitochondrial β-oxidation^28^.

Furthermore, CM patients have significantly lower levels of major catecholamine degradation metabolites, homovanillate (HVA) sulfate (FC = 10.9, B-H FDR-adjusted *P* = 7.63e-3) and vanilylmandelate (VMA) (FC = 2.2, B-H FDR-adjusted *P* = 8.17e-3) and vanillactate (VLA) (FC = 3.0, B-H FDR-adjusted *P* = 1.86e-4), compared to CM-like patients (**Fig. 4c**). HVA is the end-product of dopamine degradation while VMA is the end-product of epinephrine and norepinephrine degradation. They are synthesized through the actions of monoamine oxidase and catechol-O-methyltransferase on dopamine and norepinephrine, respectively^29^ (**Fig. 4c**). HVA and VMA are associated with catecholamine levels in the brain^30^, and their low levels suggest an impairment in catecholamine synthesis in CM patients. Catecholamines regulate membrane transport of long chain fatty acids in isolated adipocytes through beta-receptor interaction and cAMP^22^. The involvement of cAMP-dependent protein kinase suggests that the catecholamine-mediated transport process is a regulatory step in fatty acid mobilization^22^, and perturbations in catecholamine metabolism may lead dysfunctional mobilization often results in greater circulating FFA plasma concentrations^31^. The low levels of catecholamines metabolites (HVA and VMA) and high levels of thyroxine are consistent with having increased levels of circulating long chain FFAs, which suggests dysfunctional mobilization of FFAs for mitochondrial FAO in CM patients relative to CM-like patients.

### Association of steroidogenesis with coma in CM

The most significantly enriched lipid metabolism pathway in comatose patients is steroid biosynthesis (**Fig. 3d**). Notably, eight different endogenous steroids showed significantly higher levels in CM patients compared to CM-like patients (FC > 2.0, B-H FDR-adjusted *P* < 0.05). These steroids include three extracellular pregnenolone steroids (pregnenolone sulfate, pregnenediol sulfate, and pregnenetriol sulfate), three androgen steroids (16a-hydroxy DHEA 3-sulfate, andro steroid monosulfate C19H28O6S, and dehydroepiandrosterone sulfate (DHEA-S)), and two corticosteroids (cortisol and cortisone) (**Fig. 4d**).

Pregnenolone (PREG) is an immediate precursor for steroidogenesis where it is converted to dehydroandrosterone and subsequently to other androgenic steroids and corticosteroids^32,33^. These endogenous steroids are neurosteroids that are synthesized by the brain as well as the adrenal gland, gonads and T cells^34–36^, which makes their elevation in the plasma of CM patients relative to CM-like patients of significant interest. PREG levels positively correlated with edema formation and decreased neurological score in a mouse model of TBI, further supporting the contribution of some neuroactive steroids to the alterations in brain function^37^. Remarkably, the levels of multiple steroid molecules including multiple sulfated forms of PREG and its derivatives have been shown to be both elevated upon *P. falciparum* infection and positively correlated with parasitic load^38^. These steroid molecules have strong anti-inflammatory properties and impact the host immune response through inhibition of T cell proliferation and B cell immunoglobulin class switching^34,38^.

Steroids also have neuromodulatory functions and play important roles in cognitive and emotional processes, as they can act as signaling molecules and regulate various aspects of neuronal function and communication^39,40^. Our results show a consistent and significant elevation of multiple endogenous steroid molecules in CM patients with coma (**Fig. 4d**), indicating their implication in the pathophysiology of the disease. We highlight in particular the significant association between high levels of the circulating steroid hormone precursor pregnenolone sulfate (PREGS) and coma. PREGS is an endogenous neurosteroid synthesized from PREG by glial cells and neurons^41^, with cognitive and memory-enhancing, as well as antidepressant effects^42^. However, the PREGS-mediated increase in neuronal excitability, specifically through the N-methyl-D-aspartate (NMDA) and γ-aminobutyric acid A (GABA-A) receptors may become detrimental under particular conditions, leading to excitotoxic stress or convulsions^43^. Reports support a role for PREGS in coma pathogenesis, with increased levels of the neurosteroid and its neuroactive metabolite allopregnanolone in the autopsied brain tissue of cirrhotic patients who died in hepatic coma^44^. Our findings also suggest a role in the development of coma during CM.

## CONCLUDING REMARKS

Changes in plasma neurosteroid levels have been associated with neurological and psychiatric disorders^45^. However, this is the first report of a similar imbalance between comatose and non-comatose patients with severe falciparum malaria and identical brain changes on MRI. Overall, our findings demonstrate an increase of PREG levels and its downstream derivatives in comatose patients with CM, but causality remains to be established. Remarkably, PREG can modulate myelination, neuroinflammation, neurotransmission, and neuroplasticity in the brain, with potential important beneficial effects on cognition, aging, and addiction^46^. In addition, it promotes the degradation of key proteins in the innate immune signaling to suppress inflammation^42^. High levels of PREG in comatose patients could therefore be linked to a neuroprotective feedback loop triggered by the central nervous system to dampen neuroinflammation, a key mechanism of brain injury in CM^47^. Monitoring PREG levels in patients could help clinicians identify patients at risk of coma and develop more targeted treatment strategies. Our study has several limitations, including the small number of patients in each group, and a large spectrum of disease in the SNCM category, involving different organ involvement. However, this is the first study of its kin that leverages neuroimaging data to investigate the origin of coma in CM through metabolomic profiling. Further investigations are warranted to better understand the role of steroid hormones in the pathogenesis of coma in CM, with a specific focus on neuromodulation and neuroinflammation. This is particularly relevant to the current context of malaria elimination, with the global reduction in malaria transmission potentially leading to an increase in CM, as recently reported in Mozambique^48^.

## ACKNOWLEDGMENTS

We would like to thank the patients and their guardians/families for their participation in this study. We thank the Director in Charge and the clinical staff of Ispat General Hospital in Rourkela for their support and dedication, as well as the Director of the Institute of Life Sciences in Bhubaneswar for allowing us to use its Infectious Disease Biology Unit to conduct laboratory work in Rourkela.

## FUNDING

This work was supported by the National Institute of Allergy and Infectious Diseases of the National Institutes of Health under Award Numbers U19AI089676 (SCW and SM) and R21AI142472 (SCW and AH), the Medical Research Council, UK, under Award Number MR/S009450/1 (SCW and YI), and NYUAD grants ADHPG AD105 (YI and WA). HG is supported by the Science and Engineering Research Board (SERB), India, under Award Number SRG/2022/000705. The content is solely the responsibility of the authors and does not necessarily represent the official views of the funders.

## COMPETING INTERESTS

The authors declare no competing interests.

## Notes

### Competing Interest Statement

The authors have declared no competing interest.

## REFERENCES

1. Organzation, W. H. World malaria report 2022. (2022).

2. White, N. J. et al. Malaria. Lancet 383, 723–735 (2014).

3. White, N. J. Severe malaria. Malar. J. 21, 7–131 (2022).

4. Cossu, G. Therapeutic options to enhance coma arousal after traumatic brain injury: state of the art of current treatments to improve coma recovery. Br. J. Neurosurg. 28, 187–198 (2014).

5. Newton, C. R. J. C., Hien, T. T. & White, N. Cerebral malaria. J. Neurol. Neurosurg. Psychiatry 69, 433–441 (2000).

6. Medana, I. M. et al. Coma in fatal adult human malaria is not caused by cerebral oedema. Malar. J. 10, 1–14 (2011).

7. Keswani, T. et al. Pipecolic Acid, a Putative Mediator of the Encephalopathy of Cerebral Malaria and the Experimental Model of Cerebral Malaria. J. Infect. Dis. 225, 705–714 (2022).

8. Sahu, P. K. et al. Brain Magnetic Resonance Imaging Reveals Different Courses of Disease in Pediatric and Adult Cerebral Malaria. Clin. Infect. Dis. 73, E2387–E2396 (2021).

9. Mohanty, S. et al. Evidence of Brain Alterations in Noncerebral Falciparum Malaria. Clin. Infect. Dis. 75, 11–18 (2022).

10. richard oliver (dalam Zeithml., dkk 2018). Guidelines for diagnosis and treatment of malaria in India. Angew. Chemie Int. Ed. 6(11), 951–952. 2013–2015 (2011).

11. Mohanty, S. et al. Magnetic Resonance Imaging of Cerebral Malaria Patients Reveals Distinct Pathogenetic Processes in Different Parts of the Brain. mSphere 2, (2017).

12. Dehaven, C. D., Evans, A. M., Dai, H. & Lawton, K. A. Organization of GC/MS and LC/MS metabolomics data into chemical libraries. J. Cheminform. 2, (2010).

13. Evans, A. M., DeHaven, C. D., Barrett, T., Mitchell, M. & Milgram, E. Integrated, nontargeted ultrahigh performance liquid chromatography/ electrospray ionization tandem mass spectrometry platform for the identification and relative quantification of the small-molecule complement of biological systems. Anal. Chem. 81, 6656–6667 (2009).

14. Ford, L. et al. Precision of a Clinical Metabolomics Profiling Platform for Use in the Identification of Inborn Errors of Metabolism. J. Appl. Lab. Med. 5, 342–356 (2020).

15. Wishart, D. S. et al. HMDB 5.0: the Human Metabolome Database for 2022. Nucleic Acids Res. 50, D622–D631 (2022).

16. Pang, Z. et al. Using MetaboAnalyst 5.0 for LC–HRMS spectra processing, multi-omics integration and covariate adjustment of global metabolomics data. Nat. Protoc. 17, 1735–1761 (2022).

17. Goeman, J. J., Van de Geer, S., De Kort, F. & van Houwellingen, H. C. A global test for groups of genes: testing association with a clinical outcome. Bioinformatics 20, 93–99 (2004).

18. Morgan, B. L. G., Boris, G. L. & Winick, M. A Useful Correlation between Blood and Brain N-Acetylneuraminic Acid Contents. Neonatology 42, 299–303 (1982).

19. Longo, N., Frigeni, M. & Pasquali, M. Carnitine transport and fatty acid oxidation. Biochim. Biophys. Acta - Mol. Cell Res. 1863, 2422–2435 (2016).

20. Fiamoncini, J. et al. Medium-chain dicarboxylic acylcarnitines as markers of n-3 PUFA-induced peroxisomal oxidation of fatty acids. Mol. Nutr. Food Res. 59, 1573–1583 (2015).

21. Eisenhofer, G., Kopin, I. J. & Goldstein, D. S. Catecholamine Metabolism: A Contemporary View with Implications for Physiology and Medicine. Pharmacol. Rev. 56, 331–349 (2004).

22. Abumrad, N. A., Park, C. R. & Whitesell, R. R. Catecholamine activation of the membrane transport of long chain fatty acids in adipocytes is mediated by cyclic AMP and protein kinase. J. Biol. Chem. 261, 13082–13086 (1986).

23. Tyni, T., Paetau, A., Strauss, A. W., Middleton, B. & Kivelä, T. Mitochondrial Fatty Acid β-Oxidation in the Human Eye and Brain: Implications for the Retinopathy of Long-Chain 3-Hydroxyacyl-CoA Dehydrogenase Deficiency. Pediatr. Res. 2004 565 56, 744–750 (2004).

24. Khaliq, W., Andreis, D. T., Kleyman, A., Gräler, M. & Singer, M. Reductions in tyrosine levels are associated with thyroid hormone and catecholamine disturbances in sepsis. Intensive Care Med. Exp. 3, (2015).

25. Muñoz, P., Huenchuguala, S., Paris, I. & Segura-Aguilar, J. Dopamine oxidation and autophagy. Parkinsons. Dis. (2012) doi:10.1155/2012/920953.

26. Müller, M. J. & Seitz, H. J. Thyroid hormone action on intermediary metabolism - Part II: Lipid metabolism in hypo- and hyperthyroidism. Klin. Wochenschr. 62, 49–55 (1984).

27. Sayre, N. L. & Lechleiter, J. Fatty acid metabolism and thyroid hormones. Curr. trends Endocrinol. (2012).

28. Sinha, R. A., Singh, B. K. & Yen, P. M. Direct effects of thyroid hormones on hepatic lipid metabolism. Nat. Rev. Endocrinol. 14, 259 (2018).

29. Meiser, J., Weindl, D. & Hiller, K. Complexity of dopamine metabolism. Cell Commun. Signal. 11, 1–18 (2013).

30. Kopin, I. J., Bankiewicz, K. S. & Harvey-White, J. Assessment of brain dopamine metabolism from plasma HVA and MHPG during debrisoquin treatment: validation in monkeys treated with MPTP. Neuropsychopharmacology 1, 119–125 (1988).

31. Beard, J. K. & Yates, D. T. Function and dysfunction of fatty acid mobilization: a review. Diabesity 5, (2019).

32. Miller, W. L. & Bose, H. S. Early steps in steroidogenesis: Intracellular cholesterol trafficking. Journal of Lipid Research vol. 52 2111–2135 (2011).

33. Wassif, W. S. & Ross, A. R. Steroid Metabolism and Excretion in Anorexia Nervosa. Anorexia 92, 125–140 (2013).

34. Mahata, B. et al. Single-cell RNA sequencing reveals T helper cells synthesizing steroids De Novo to contribute to immune homeostasis. Cell Rep. 7, 1130–1142 (2014).

35. Mahata, B. et al. Tumors induce de novo steroid biosynthesis in T cells to evade immunity. Nat. Commun. 11, 1–15 (2020).

36. Pearce, E. J. et al. Th2 response polarization during infection with die helminth parasite Schistosoma mansoni. Immunol. Rev. 201, 117–126 (2004).

37. Lopez-Rodriguez, A. B. et al. Profiling neuroactive steroid levels after traumatic brain injury in male mice. Endocrinology 157, 3983–3993 (2016).

38. Abdrabou, W. et al. Metabolome modulation of the host adaptive immunity in human malaria. Nat. Metab. 2021 37 3, 1001–1016 (2021).

39. Frye, M., Jaffrey, S. R., Pan, T., Rechavi, G. & Suzuki, T. RNA modifications: what have we learned and where are we headed? Nat. Rev. Genet. 17, 365–372 (2016).

40. Remage-Healey, L. Frank Beach Award Winner: Steroids as Neuromodulators of Brain Circuits and Behavior. Horm. Behav. 66, 552 (2014).

41. Agís-Balboa, R. C. et al. Characterization of brain neurons that express enzymes mediating neurosteroid biosynthesis. Proc. Natl. Acad. Sci. U. S. A. 103, 14602–14607 (2006).

42. Murugan, S., Jakka, P., Namani, S., Mujumdar, V. & Radhakrishnan, G. The neurosteroid pregnenolone promotes degradation of key proteins in the innate immune signaling to suppress inflammation. J. Biol. Chem. 294, 4596–4607 (2019).

43. Williamson, J., Mtchedlishvili, Z. & Kapur, J. Characterization of the convulsant action of pregnenolone sulfate. Neuropharmacology 46, 856–864 (2004).

44. Ahboucha, S., Pomier-Layrargues, G., Mamer, O. & Butterworth, R. F. Increased levels of pregnenolone and its neuroactive metabolite allopregnanolone in autopsied brain tissue from cirrhotic patients who died in hepatic coma. Neurochem. Int. 49, 372–378 (2006).

45. Ratner, M. H., Kumaresan, V. & Farb, D. H. Neurosteroid Actions in Memory and Neurologic/Neuropsychiatric Disorders. Front. Endocrinol. (Lausanne). 10, 169 (2019).

46. Lin, Y. C., Cheung, G., Espinoza, N. & Papadopoulos, V. Function, regulation, and pharmacological effects of pregnenolone in the central nervous system. Curr. Opin. Endocr. Metab. Res. 22, 100310 (2022).

47. Idro, R., Marsh, K., John, C. C. & Newton, C. R. J. Cerebral malaria: mechanisms of brain injury and strategies for improved neurocognitive outcome. Pediatr. Res. 68, 267–274 (2010).

48. Guinovart, C. et al. The epidemiology of severe malaria at Manhiça District Hospital, Mozambique: a retrospective analysis of 20 years of malaria admissions surveillance data. Lancet Glob. Heal. 10, e873–e881 (2022).

